# Comparison of hemodynamic response functions obtained from resting-state functional MRI and invasive electrophysiological recordings in rats

**DOI:** 10.1101/2023.02.27.530359

**Authors:** D Rangaprakash, Olivier David, Robert L Barry, Gopikrishna Deshpande

## Abstract

Resting-state functional MRI (rs-fMRI) is a popular technology that has enriched our understanding of brain and spinal cord functioning, including how different regions communicate (connectivity). But fMRI is an indirect measure of neural activity capturing blood hemodynamics. The hemodynamic response function (HRF) interfaces between the unmeasured neural activity and measured fMRI time series. The HRF is variable across brain regions and individuals, and is modulated by non-neural factors. Ignoring this HRF variability causes errors in FC estimates. Hence, it is crucial to reliably estimate the HRF from rs-fMRI data. Robust techniques have emerged to estimate the HRF from fMRI time series. Although such techniques have been validated non-invasively using simulated and empirical fMRI data, thorough invasive validation using simultaneous electrophysiological recordings, the gold standard, has been elusive. This report addresses this gap in the literature by comparing HRFs derived from invasive intracranial electroencephalogram recordings with HRFs estimated from simultaneously acquired fMRI data in six epileptic rats. We found that the HRF shape parameters (HRF amplitude, latency and width) were not significantly different (*p*>0.05) between ground truth and estimated HRFs. In the single pathological region, the HRF width was marginally significantly different (*p*=0.03). Our study provides preliminary invasive validation for the efficacy of the HRF estimation technique in reliably estimating the HRF non-invasively from rs-fMRI data directly. This has a notable impact on rs-fMRI connectivity studies, and we recommend that HRF deconvolution be performed to minimize HRF variability and improve connectivity estimates.

## 1. Introduction

Blood oxygenation level-dependent (BOLD) functional magnetic resonance imaging (fMRI) has been a powerful technology for non-invasively measuring central nervous system (CNS) function [1]. Unlike electroencephalogram (EEG), which directly measures neuronal electrical activity, fMRI only indirectly measures neural activity through local blood oxygenation variations [2] (**Figure 1**). Although oxygen demand is proportional to neural activity, it is also modulated by dilation/constriction of blood vessels that are influenced by several non-neural factors and are thus difficult to delineate [3] (neurovascular coupling). These non-neural factors influence the BOLD signal (variable density/size of vessels, caffeine/alcohol/lipid intake, magnetic susceptibilities, hematocrit, partial volume imaging, respiration/pulse differences [1] [4] [5] [6] [7] [8] [9] [10]). This combination of factors interfacing neural activity to BOLD fMRI is the hemodynamic response function (HRF) [11] [12]. It changes across the brain and individuals [4] [5] [13], since the non-neural factors are variable too.

**Figure 1.**
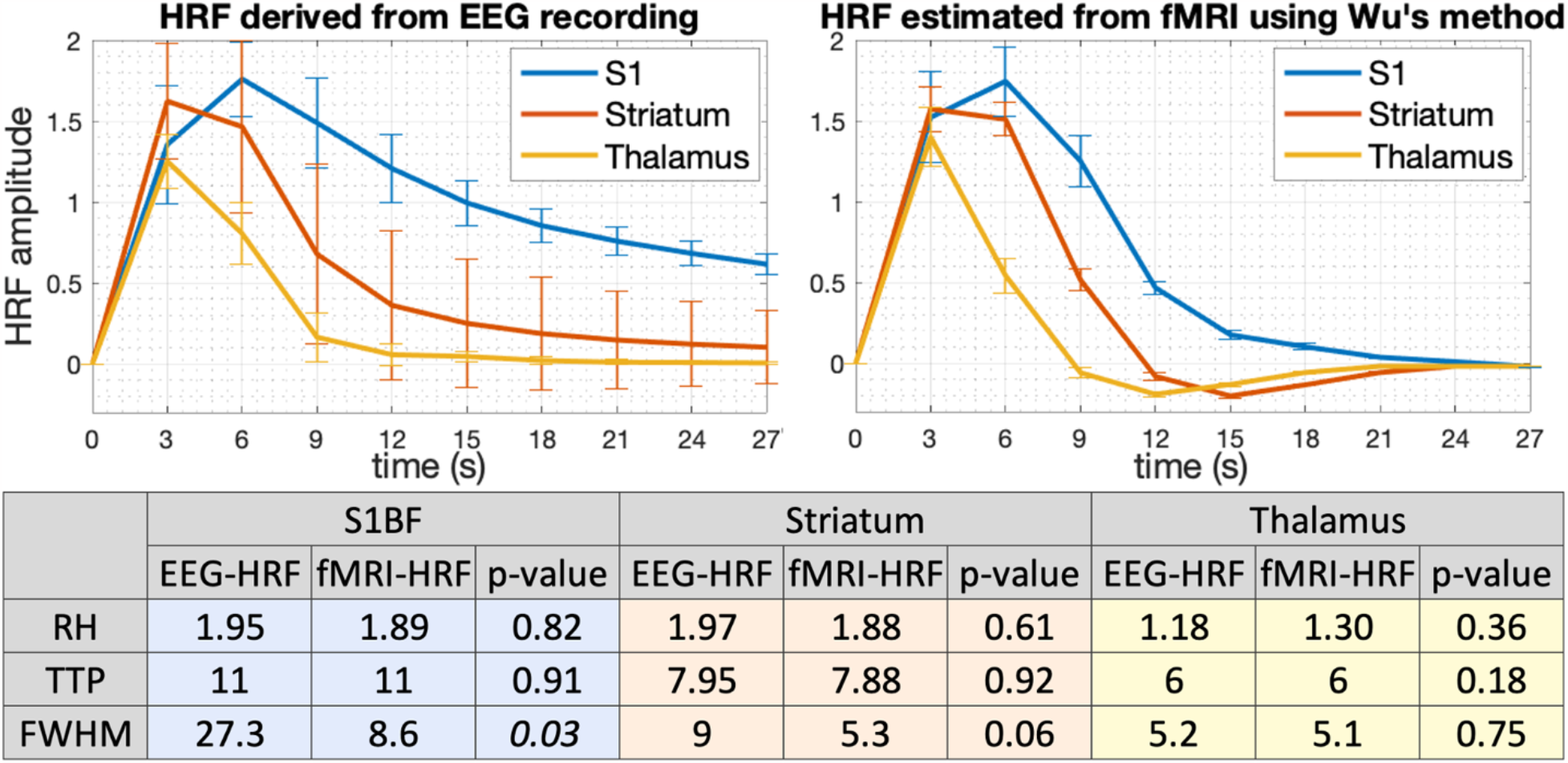
Plots of ground truth iEEG-derived HRF (top-left) vs. Wu et al.’s estimated HRF (top-right). HRF shape parameters were not significantly different between ground truth and estimated HRFs (p>0.05) except for FWHM in one pathological region (p=0.03). The table at the bottom shows median values along with p-values for comparison between ground truth and estimated HRFs (paired t-test was performed for RH, and Wilcoxon sign-rank test was performed for TTP and FWHM). HRF is positive in amplitude; RH is unitless (only relative differences matter); TTP and FWHM are in seconds; error bars denote standard deviations.

It has been found that this HRF variability (HRFv) is undesirable in fMRI because it confounds resting-state fMRI (rs-fMRI) connectivity estimates. Specifically, ignoring HRFv results in about 15% error in functional connectivity (FC) estimates in both the brain and spinal cord [14] [15], causing the detection of false positive and false negative connections. Effective connectivity modeling is also viable only after minimizing HRFv [16] [17]. Hence, estimating the HRF and minimizing HRFv is critical to the vast rs-fMRI field.

HRFs estimated from task fMRI data [18] [19] [20] [21] [22] [23] [24] are not directly applicable to rs-fMRI data because a specific task does not activate the entire brain uniformly [25] (HRFs are biased to activated voxels). Even in activated voxels, the BOLD time series could be non-uniform [26], leading to biologically implausible HRF estimates [25]. Further, evidence suggests that spontaneous fluctuations in rs-fMRI can be observed at frequencies not supported by the canonical HRF [27] [28] [29] [30] [31]. Therefore, it is advisable to directly estimate the HRF from rs-fMRI data to minimize the HRF confound in rs-fMRI connectivity modeling. The HRF cannot be directly measured non-invasively with rs-fMRI, and it may not be possible to estimate HRF latency from a hypercapnic challenge [32] due to the unavailability of a breath-hold fMRI scan or holding breath not being feasible in pathological populations [33].

However, critical technical advancements such as point-process theory [34] [35] have now provided us with deconvolution techniques that can perform HRF estimation with rs-fMRI alone [36] [37] [38] [39]. They have been validated [34] [36] [14] and applied in recent studies [40] [41] [42] [43] [44] [45] [46]. These deconvolution techniques estimate two unknowns (latent neural activity and the HRF) using one input (fMRI time series), which is possible in a least-squares sense because we know the biophysics and natural limits of HRF and fMRI, which constrain the solution space. Although a few excellent rs-fMRI HRF deconvolution techniques are available [47] [37] [38] [39], here we specifically focus on Wu et al.’s method [36]. Extending our results to the other techniques is within the scope of future work.

Wu et al.’s method [36] considers rs-fMRI as event-related time series with events modeled as point-processes [48]. It then estimates the least-squares best fit HRF from the fMRI time series at identified events. Latent neural time series is estimated from fMRI and HRF using Wiener deconvolution [49]. This technique is validated non-invasively using simulations and empirical fMRI data [34] [36] [14]; it is well-recognized (∼250 citations) and has been utilized in several recent papers (e.g., [14] [40] [41] [42] [43] [44] [45] [46]). However, invasive validation using independent electrophysiological recordings, the gold standard, has been elusive. We addressed this gap by utilizing simultaneous invasive intracranial EEG (iEEG) recordings and fMRI data acquired in six epileptic rats (thorough data acquisition details are in [50] [51]). We compared the HRFs derived from iEEG with those estimated from fMRI.

## 2. Materials and methods

### 2.1 Data acquisition

All the data used by us were previously acquired for a different project, and herein we performed further processing and secondary analyses to achieve our goals. Complete details of animal preparation, experimental setup, data acquisition and data denoising/preprocessing can be found in [50]. We briefly describe it here for the benefit of the readers.

Data were acquired from Genetic Absence Epilepsy Rats from Strasbourg (GAERS) [52]. These rats exhibited spontaneous spike and wave discharges (SWDs) lasting about 20s and repeating every minute at rest. These epileptic seizures stemmed from an epileptic focus and spread to other brain regions [52]. SWDs originate in perioral regions of S1 (first somatosensory cortex, S1BF) [53] [54]; that is, this region is pathogenic in these rats.

Animal care/experiments were conducted in accordance with the European Community Council Directive of 24 November 1986 (86/609/EEC). They were approved by the animal experimentation Ethical Committee at the Université Joseph Fourier, Grenoble, France (protocol number 88–06). Six male adult GAERS were included (281±56g). Spontaneous SWDs were measured with iEEG during MRI data acquisition. Three carbon electrodes situated in the skull near the midline were used (frontal, parietal and occipital) for iEEG data acquisition from a digital acquisition system (Cambridge Electronic Design); sampling rate = 2kHz, analog filter passband = 1–90Hz. Two reference electrodes were placed in the occipital and nasal bones. FMRI and iEEG temporal co-registration was done by the iEEG acquisition software using the MRI system’s trigger signal during each volume acquisition. MRI data were acquired in a horizontal-bore 2.35 Tesla scanner (Bruker Spectrospin) having actively shielded gradient coils (Magnex Scientific). A linear volume transmit coil and an actively decoupled surface receive coil were used. A gradient-echo echo planar imaging sequence was used for fMRI: two shots, TR/TE = 3000/20 ms, flip angle = 90°, voxel size = 0.73×0.73×1.5 mm^3^, 15 slices, whole brain coverage. Scan duration ranged from 30 to 150 minutes across rats. T1-weighted anatomical data were acquired with a 3D-MDEFT sequence (modified driven equilibrium Fourier transform) [55]: TR/TE = 15/5 ms, inversion time = 605 ms, flip angle = 22°, voxel size = 0.33×0.33×0.33 mm^3^.

SPM 5 [56] was used for data processing and quality assurance. Standard spatial preprocessing was performed, including realignment, co-registration and smoothing. Three ROIs were identified [50]: primary somatosensory cortex barrel field (S1BF), thalamus and striatum. Activations were found in S1BF and thalamus, whereas deactivations were found in the striatum [50]. These three regions were selected because (1) they were consistently activated across different sessions and rats; (2) they exhibited a wide range of hemodynamic responses, which is helpful for rigorous validation of our estimated HRFs; and (3) these regions align well with our current understanding of SWDs. Mean time series from these three regions were used in subsequent analysis. Please refer to [50] [51] for complete details of data acquisition and preprocessing.

### 2.2 Data processing and analyses

All fMRI data processing presented here, as well as the comparison of HRFs, was carried out exclusively for this study. As described before, HRFs were estimated from fMRI time series separately in each ROI of each rat using Wu et al.’s method [36]. It considers rs-fMRI as event-related time series; events are modeled as point-processes (threshold: mean+1SD) [48]. It then estimates the best fit HRF at identified events in the fMRI time series in a least-squares sense. Latent neural time series is then estimated from HRF and fMRI using Wiener deconvolution [49]. A temporal mask with aggregate motion (framewise displacement) <0.3mm is used to avoid false events driven by motion [35]. The HRF shape can be characterized by three parameters (see Figure 1 in [14]) – response height (RH or HRF amplitude), time-to-peak (TTP or HRF latency) and full-width-at-half-max (FWHM or HRF width).

The procedure for deriving the HRFs from iEEG is explained in detail in David et al. [50] (the same iEEG-derived HRFs from that study were used here, with inverted HRFs rendered as positive). Briefly, the ground-truth HRF of each ROI in each rat was obtained using Wiener deconvolution within a biologically-constrained biophysical model of brain hemodynamics [57] [58], with the iEEG time series as the input, fMRI time series as the output and the HRF as the transfer function between the two that was derived. HRF parameters (RH, TTP and FWHM) were then quantified from the derived HRFs.

The mechanism of how the Wu et al. method estimates the HRF, how it relates to the underlying neurovascular physiology, and what the estimated events correspond to has been addressed in previously published studies [36] [34] [59]. The current focused technical note specifically addresses the question of whether, and how well, the HRFs estimated from Wu et al.’s method correspond to ground-truth HRFs, in the specific case of epileptic rats. We statistically compared the HRF shape parameters estimated from fMRI with the corresponding iEEG-derived HRF parameters. We compared the HRF shape parameters (see Figure 1 in [14]) – RH, TTP and FWHM – in all three regions across the six rats. A paired t-test was performed for RH (continuous variable), and a Wilcoxon sign-rank test was performed for TTP and FWHM (discrete variables, with the smallest unit being one sample or TR). The only variability that mattered for the statistics was the difference between estimated and ground truth HRFs within a given region of a given rat, because paired tests are unaffected by between-sample variability.

## 3. Results

Our three ROIs were S1BF, striatum and thalamus. Since S1BF was the pathological region (epileptic focus), an abnormally slow HRF was observed compared to the striatum and thalamus (**Figure 1**). This is understood to be due to the suppression of cerebral blood flow autoregulation mechanisms modulating vasodilatation. Please refer to David et al. [50] for descriptions and other analyses related to the iEEG data used here.

We found an excellent match between the HRFs (**Figure 1**). RH and TTP were not significantly different between estimated and ground truth HRFs in all regions (*p*>0.05), and FWHM was not different except in S1BF. In S1BF, FWHM was significantly different (*p*=0.03) without multiple comparisons correction, and would not have been significantly different with either FDR or Bonferroni correction. The original study [50] describes S1BF as pathological with abnormal HRF. Despite this, the relative HRF amplitude and timing information were preserved. Overall, except for the FWHM misspecification in a pathological region, the HRF shape estimated from Wu et al.’s technique was not significantly different from the ground truth HRF shape (**Figure 2**).

**Figure 2.**
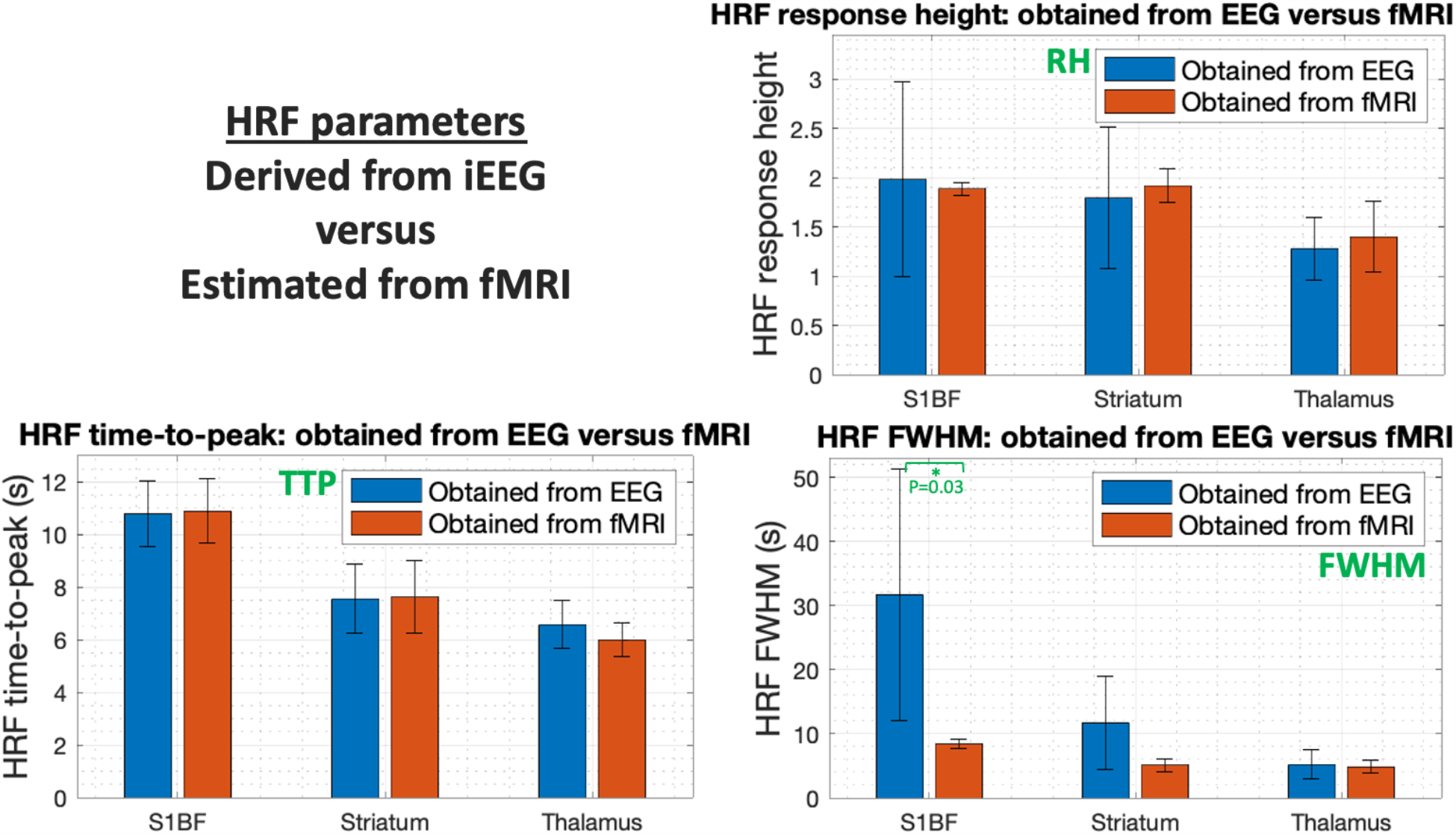
Bar graphs comparing HRF shape parameters of iEEG-derived ground truth HRF and fMRI-estimated HRF in the three regions of interest. All parameters in all regions were not significantly different except FWHM in the S1BF region, which was different with p=0.03. Error bars denote standard deviations.

## 4. Discussion

Our goal was to compare HRFs estimated non-invasively from fMRI data (without knowledge of the neural input) with ground-truth HRFs derived invasively using iEEG (with knowledge of the neural input). We tested this in the specific case of three ROIs in six epileptic rats. We found that estimated and ground-truth HRFs were not significantly different, with the only exception of one of the shape parameters in one pathological region. Even in this region (S1BF), the HRF amplitude (RH) and timing (TTP) were estimated accurately, but the HRF width (FWHM) was underestimated. Taken together, we concluded that our results provide preliminary invasive validation for Wu et al.’s technique’s ability to estimate the HRF from fMRI data in non-pathological regions in rats.

We now discuss a couple of criticisms that could be raised about the results. One argument is that Wu et al.’s method performed well on this dataset because the larger amplitude of events due to pathology likely boosted the results artificially. However, that is not the case because there was only one pathological region (S1BF), and the other two regions were found to be normal [50]. The algorithm did not perform significantly better in S1BF – rather the ground truth and estimated HRFs were found to be not significantly different in the non-pathological regions with even larger p-values. Moreover, HRF shape is characterized not only by its amplitude but also by TTP and FWHM. The Wu et al. method is data-driven and is blind to where the data comes from, so there is no inherent bias in the method arising from *a priori* knowledge of pathology.

Another question of interest is what accounts for the larger variability in the ground-truth HRFs (Figure 1) as compared to Wu’s et al.’s estimated HRFs. Several speculative statements can be made about why estimated HRFs had lesser variability across regions and rats compared to ground-truth HRFs. One is that the Wu et al. method estimates the HRF within tight biological constraints of typical HRFs based on the balloon model [60] and the double-gamma HRF fit [4], which likely reduces the degrees of freedom of the estimated HRF thus lowering variability. Deviating from the ground truth by having lower variance is preferable to deviating with too much variance; the former would minimally impact deconvolved latent neural time series used in further post-processing while the latter would significantly meddle with subsequent time series analyses. Nevertheless, these statements are speculative, and we believe that it is infeasible to derive a definitive answer with our available data.

Our study has a few caveats and limitations. (i) Whether our results extend to humans in both healthy and pathological cases remains to be explored, but it is not straightforward to test because of the invasive nature of the experimental setup and the necessity for simultaneous iEEG and fMRI acquisition. (ii) Although the rats were at rest during the experiments and not involved in any active task, these were pathological rats having seizures (switching between ictal and interictal states); hence, extending our results to human rs-fMRI requires further assessment. (iii) Our sample size was small (N=6). Yet, we believe that our sample provided preliminary evidence with reasonable confidence. Further, when comparing conditions within subjects, the sample size requirements are rather different. For example, Crowe et al. published a study [61] with just two monkeys since they did robust within-subject comparisons. The same holds true for David et al.’s original paper using this data [50] where they compared the ground truth effective connectivity with that obtained from dynamic causal modeling. (iv) The impact of fMRI temporal resolution on our results is also unclear because our validation with low temporal resolution data (TR=3s) may or may not extend to higher resolutions (TR<1s) that are not uncommon today.

Despite these limitations, our results are nevertheless encouraging. The HRF confound is a key issue that is difficult to ignore, and rs-fMRI connectivity is an expanding field of research with dozens of papers published weekly. Given this context, our results are significant because most rs-fMRI studies ignore HRFv and do not perform deconvolution, with one of the reasons being that the HRF estimation techniques lack validation using ground truth obtained from invasive procedures. Our study entrusts a level of confidence in the Wu et al. technique and its ability to estimate the HRF reliably, and provides motivation to conduct further invasive validation studies under various scenarios in humans. Such efforts are also important for other HRF estimation techniques.

The study reported herein is not the first study to attempt invasive validation of Wu et al.’s technique. A recent study from the same group [59] used co-localized intrinsic optical signals (IOS) and local field potentials (LFPs) in a rat to compare the HRF estimated using Wu et al.’s technique from IOS signals with the HRF derived from the envelope of the LFP signal. They found a high correlation between the two HRFs (*R*=0.82±0.04). However, our study directly used fMRI data instead of IOS signals, which is important because each modality has its stereotypical noise profiles and idiosyncratic processing pipelines that can make a difference to the outcomes. Hence, it is reaffirming that our results validated the Wu et al. method using fMRI data, corroborating what their study validated with IOS signals. Our study also included six rats instead of one, and three ROIs instead of one. The conclusion from our study supports the conclusions from theirs.

Taken together, our findings support the notion that it is possible to accurately estimate the HRF non-invasively from rs-fMRI data, at least in non-pathological regions in rats. Further research must confirm this in humans under both healthy and pathological cases. We encourage researchers to account for HRFv and minimize the HRF confound in their rs-fMRI connectivity studies. We recommend performing HRF deconvolution and utilizing the estimated latent neural time series in subsequent analyses instead of directly using the BOLD time series.

### CRediT (Contributor Roles Taxonomy) – author contributions

**D Rangaprakash**: Conceptualization, Methodology, Data Analysis, Investigation, Visualization, Writing - Original Draft, Reviewing and Editing. **Olivier David**: Data Acquisition, Investigation, Writing - Reviewing and Editing. **Robert L Barry**: Funding Acquisition, Project Administration, Investigation, Writing - Reviewing and Editing, Supervision. **Gopikrishna Deshpande**: Funding Acquisition, Project Administration, Conceptualization, Methodology, Investigation, Writing - Reviewing and Editing, Supervision.

## Data availability statement

Study data can be made available upon reasonable request to the corresponding author.

## Acknowledgments

Secondary data analyses were supported in part by the National Institutes of Health (NIH) through grants R01EB027779 and R21EB031211 (R.L.B.). The content is solely the responsibility of the authors and does not necessarily represent the views of the NIH. The funders had no role in study design, data collection and analysis, decision to publish, or preparation of the manuscript.

